# Modelling co-translational dimerisation for programmable nonlinearity in synthetic biology

**DOI:** 10.1101/2020.07.10.196667

**Authors:** Ruud Stoof, Ángel Goñi-Moreno

## Abstract

Nonlinearity plays a fundamental role in the performance of both natural and synthetic biological networks. Key functional motifs in living microbial systems, such as the emergence of bistability or oscillations, rely on nonlinear molecular dynamics. Despite its core importance, the rational design of nonlinearity remains an unmet challenge. This is largely due to a lack of mathematical modelling that accounts for the mechanistic basics of nonlinearity. We introduce a model for gene regulatory circuits that explicitly simulates protein dimerization—a well-known source of nonlinear dynamics. Specifically, our approach focusses on modelling co-translational dimerization: the formation of protein dimers during—and not after—translation. This is in contrast to the prevailing assumption that dimer generation is only viable between freely diffusing monomers (i.e., post-translational dimerization). We provide a method for fine-tuning nonlinearity on demand by balancing the impact of co- versus post-translational dimerization. Furthermore, we suggest design rules, such as protein length or physical separation between genes, that may be used to adjust dimerization dynamics in-vivo. The design, build and test of genetic circuits with on-demand nonlinear dynamics will greatly improve the programmability of synthetic biological systems.

## 1 Introduction

Synthetic biology [1, 2, 3] is a growing field of research that uses engineering principles to design and implement human-defined computations in living cells. The development of complex mathematical modelling techniques [4, 5], along with the advances in DNA synthesis and assembly [6], allows for the making of new-to-Nature networks of regulatory proteins—the so-called genetic circuits—with increasing information processing abilities. Relatively simple models of computation, like combinatorial [7] and sequential logic [8], have been successfully implemented in organisms such as bacteria [9], yeasts [10] or even mammalian cells [11]. Examples of these genetic circuits include logic gates [12, 13], multiplexers [14], half-adders [15], counters [16] and memories [17]—even more complex processes such as analogue [18, 19] and distributed computations [20, 21, 22]. As well as generating fundamental insights into the workings of living systems, this (and many more) engineered systems find a broad range of applications in biotechnology and bioengineering [23, 24]. Nevertheless, it could be argued that the field is in its infancy, since the vast differences between traditional silicon-based hardware and biological substrate suggest there are powerful models of computation yet to be developed [25]. This is, the way synthetic constructs process information cannot get even close to that of natural biological systems.

Mathematical modelling, an essential component of synthetic biology, aids the design of increasingly complex genetic circuits by providing predictions of their dynamical performance [26, 27]. By modelling the dynamics of gene expression (i.e. mechanistic steps going from genes to proteins), designs can be improved with information such as, for instance, which DNA components are predicted [28] to perform better for a specific target function. This is, models answer questions that are difficult, if not at all impossible, to answer otherwise. However, the predictive power of models depends on the detail of the description of the physical processes they represent. There is one such processes, that, despite being of fundamental importance for genetic circuits and biological computation, has received relatively little attention to this end: the mechanistic understanding of nonlinear dynamics [29, 30, 31, 32]. Specifically, due to its relevance for genetic circuits, the nonlinearity that emerges from the dimerisation of regulatory proteins [33]. Upon gene expression, most resulting proteins are *monomers* that need to interact to form *dimers* (or higher-order *oligomers);* it is only the protein in its final from that is active. Furthermore, dimerisation is intertwined with other sources of nonlinearity [34], such as regulator-promoter interplay [35] (known as *bursting*) In the early days of synthetic biology, two landmark papers demonstrated that such nonlinear dynamics are at the core of relatively simple circuits. The first one implemented a “toggle switch” [36]; a genetic circuit able to switch between two states according to external signals. The second one implemented what was called the “repressilator” [37]; an oscillator based on a circuit of gene transcription repressors. Mathematical models of these two circuits show bistability and reliable oscillations (respectively) only if they account for nonlinear dynamics.

A non-trivial issue is that protein monomers have to *meet* in order for dimerisation to happen. But, where do they meet? Nearly thirty years ago, a paper titled “[…] it takes two to tango but who decides […] on the venue?” [38] fired this question explicitly; and soon after, another paper found experimental evidence for what they called *co-translational* dimerisation [39]. The latter work established a difference between co-translational, which takes place *during* translation (from RNA into proteins), and post-translational dimerisation pathways. However, current mathematical modelling accounts for the dimerisation of proteins using abstract parameters without physical description. While Michaelis-Menten equations, for instance, use integer values for a physical parameter, Hill equations allow for unphysical non-integer values. Although somewhat effective, these modelling approaches abstract away dimerisation into a black box, thus hiding it from the design process. The lacking description of the molecular details of nonlinearity limits the rational design of such behaviour. To overcome this limitation is the target of the present research.

## 2 Results and Discussion

### 2.1 A model for co-translational dimerization

Transcription (of a gene into RNA) and translation (of RNA into proteins) rarely generate fully functional transcriptional regulators. Rather, resulting proteins—or “monomers”—need to interact with others in order to form “dimers” (1A); for example, the repressor protein TetR, which is extensively used in synthetic biology, is a dimer. This suggests that any mathematical model that aims at simulating the dynamics of such a molecule (or that of a genetic circuit regulated by it) must account for the dimerisation of the partially formed regulators generated after translation in order to result in robust predictions.

Although typical modelling frameworks as, for instance, Michaelis-Menten and Hill equations, already account for dimerisation, matching this to its specific molecular mechanisms and features (in contrast to using abstract cooperativity values) is still an overarching challenge. Our model adds a detailed mechanism of translation, in which ribosomes bind RNA molecules in an asynchronous fashion; as a result, some ribosomes start translating very close to each other [40]. In this scenario, two partially-formed monomers come into contact and dimerisation starts, which has been termed co-translational dimerisation (1B) [39].When ribosomes are at a long-enough distance, dimerisation will take place in the cytoplasm (i.e., post-translational dimerisation), where both monomers must meet before building a dimer. The model presented here accounts for these dynamics, which, in turn, shape the nonlinearity features (in final dimer formation) of the genetic circuit at stake.

In order to describe co-translational dimerisation rates, the model firstly assume that the arrival of a ribosome to the RNA is captured by a Poisson process. The distribution of time between binding events (*P*(Δ*t*)) is then given by the following equation:

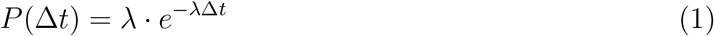

where λ is the average binding rate of a ribosome to the RNA (i.e., when translation initiates), and Δ*t* represents the time between such binding events. This time distribution, along with experimentally obtained translation rate of ca. 10 *AA/s* [41, 42] (*AA* = amino acids), was used to calculate the distance between two consecutive ribosomes (Δ*x*) and the length difference of the partially-translated monomers being generated by them (Δ*r*).

To calculate the fraction of dimerisation that takes place *during* translation (versus *after* translation), the model defines the area where partially formed monomers can interact (Figure 1B). While monomers are being formed, they are bound to the ribosome in one end and moving within a sphere around the ribosome, with a radius based on the current length, with the other end. If two monomers are long enough, the volume where the spheres overlap shows where the monomers can interact. In this state, co-translational dimerisation may start. The relative volume for these overlapping spheres (with distance between origins Δ*x*, and radii *r* and *r* + Δ*r*) is:

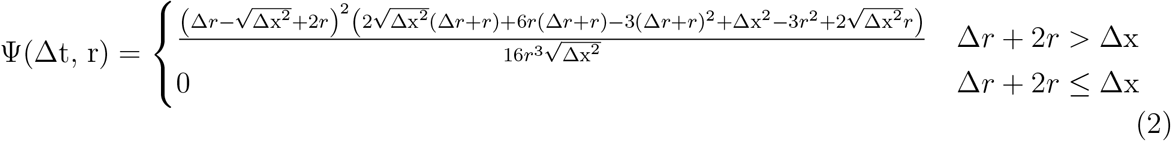

where the output is a relative value going from 0 (no overlap at all) to 1 (complete overlap of partially formed monomers), Δ*x* is the distance between two ribosomes, and *r* is the current length of the least transcribed monomer (see Methods for model derivation). The creation of dimers at each specific time difference and at specific protein length is assumed to obey Michealis-Menten dynamics. Therefore, the calculation of the fraction of dimerisation during translation (*α*) is given by the following equation:

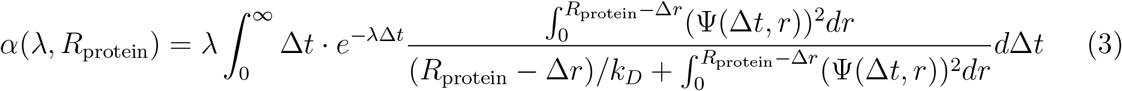

where *R*_protein_ is the final length of the protein and *k_D_* is a rate termed *reaction parameter* that accounts for the likeliness of two amino acid chains to dimerise when they co-exist in the same volume.

**Figure 1:**
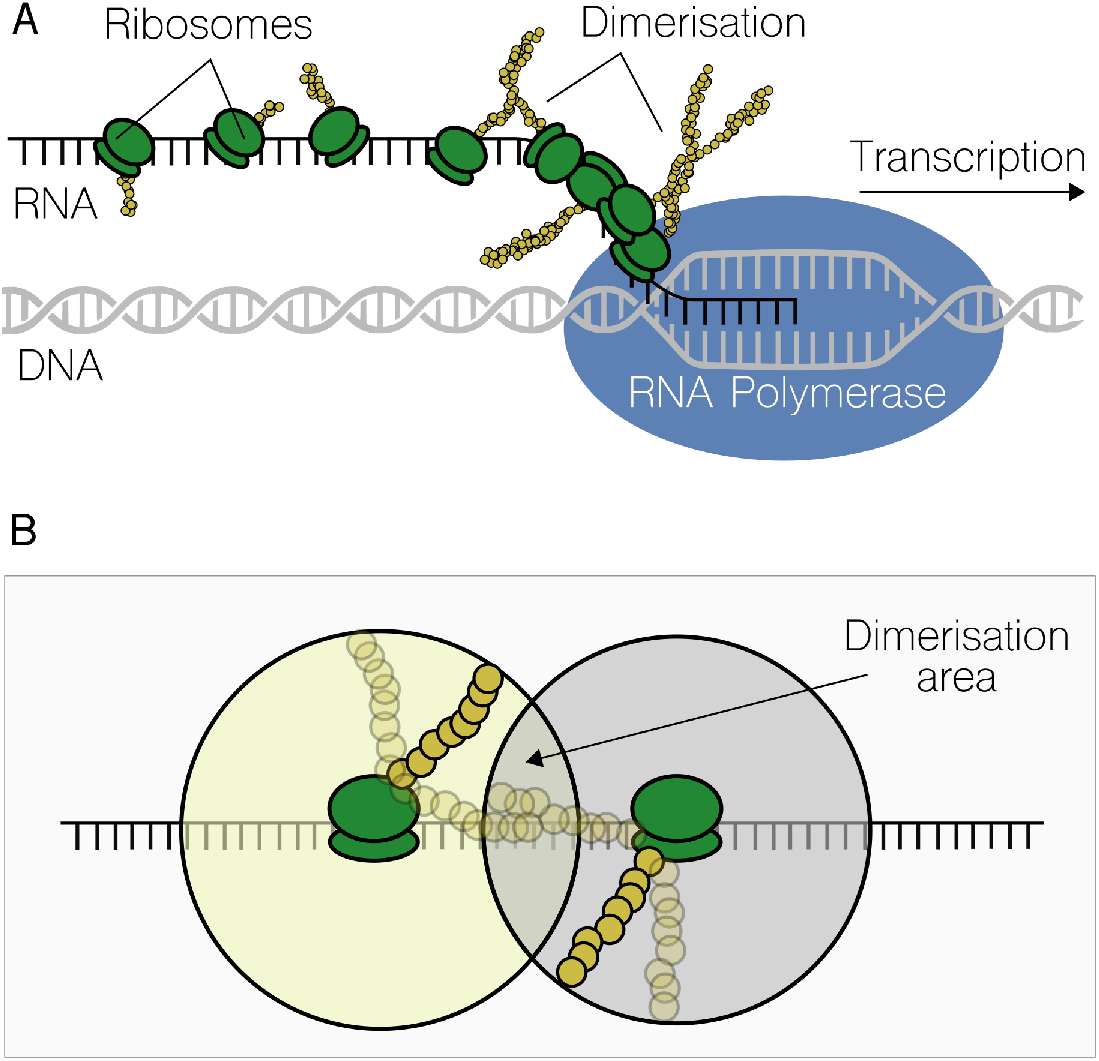
Co-translational dimerisation. **A.** Upon DNA transcription by RNA polymerase, ribosomes bind the resulting RNA to translate it into proteins. There is more than one translation processes at any one time, and ribosomes go along the RNA at different speeds, leading to the appearance of *traffic jams*. Our model simulates the process by which, when distance among ribosomes is short, partially formed monomers (represented in the figure by chains of yellow circles) dimerise with other partially formed monomers as they are being translated. **B**) Detail of the translation-mediated dimerisation area. The extension of this region depends on several physical features, as the length of the protein to be translated or the distance between ribosomes; these constraints will affect nonlinearities due to protein dimerisation.

Figure 2 shows the resulting fraction of co-translated dimerisation (*α*) in relation to key model parameters (Figure 2A): the binding rate of the ribosomes (λ), the length of the monomers (measured in amino acids) and the reaction parameter (*k_D_*). These results are shown for two different values of the reaction parameter (dimensionless), 1 (Figure 2B) and 10 (Figure 2C), at experimentally obtained values for the binding rate [43], λ ≈ 0.1*s*^-^1. As observed here, *α* is minimum (from 0 to ≈0.2) at lower values of λ and protein length; although λ is the limiting rate here, since at very low values the protein length does not make a difference. This suggests that a very low binding rate (or low ribosome availability) would result in having no co-translational dimerisation—or small values which could be neglected. However, as soon as λ increases, *α* gains importance, also amplified by increasing the reaction parameter *k_D_*. This is because the more λ increases, the more ribosomes will bind to the RNA within a given time interval; as a consequence, ribosomes will be physically closer while translating and partially-formed monomers will tend to overlap more (i.e., higher values at equation 2).

**Figure 2:**
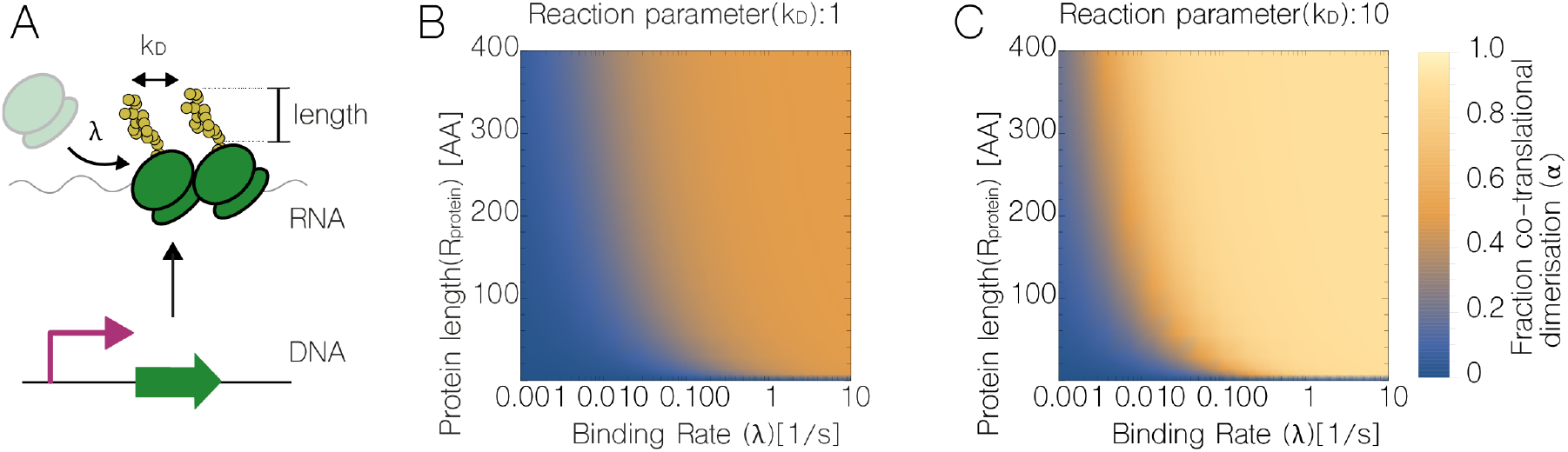
Mechanistic principles of co-versus post-translational dimerisation dynamics. **A.** Diagram that shows the modelling parameters affecting dimerisation types: ribosome binding (λ), the rate with which two partially-formed monomers interact (*k_D_*) and protein length. **B.** At a rate where monomers weakly interact (*k_D_* = 1, dimensionless) the fraction of co-translational dimerisation (i.e., total dimerisation minus dimerisation in the cellular cytosol), *α*, as described by equation 3, responds primarily to ribosome binding: if binding increases (x axis), *α* also increases (colour map). **C.** The pattern in *α* when monomers strongly interact (*k_D_* = 10) is similar, but with a (much) sharper transition from lower to higher values.

The results shown in Figure 2 suggest there is a fragile equilibrium in the fraction of co-translational vs. post-translational (i.e., in the cytosol) dimerisation, which, in turn, would impact on the nonlinear dynamics of the system. Therefore, different values of *α* will modify the performance of genetic circuits that build on nonlinear reactions to achieve optimal behaviour. In what follows, we analyse how *α* may be used to alter oscillations and bistability.

### 2.2 Case study: programmable nonlinearity in a genetic oscillator

A general requirement in order to obtain oscillations from a biological system is that its kinetic machinery must be “sufficiently nonlinear” [44]. Here, we build a mathematical model for a theoretical three-component genetic oscillator, based on the “repressilator” equations [45]. The model accounts for the new parameter that represents the fraction of dimerisation that takes place during translation (*α*), and simulates how this parameter can, by itself, modify the nonlinearity of the system.

This genetic circuit is formed by a ring of repressors (proteins that negatively regulate a promoter), where each one regulates its successor (Figure 3A). As a result, the concentration of all three proteins within the circuit oscillates in time. The system of Ordinary Differential Equations (ODEs) that represents the oscillator is described as follows. Firstly, the ODE that calculates the concentration of monomeric repressors is:

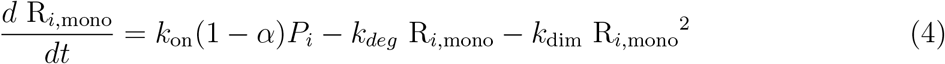

where *i* ∈ 1, 2, 3 is a specific repressor protein, *R_mono_* is the amount of repressor monomer in the cytosol (i.e., those monomeric regulators that did not dimerise during translation), *k_on_* is the expression rate of monomers when the promoter P is not repressed (thus fully active), *k_deg_* represents the degradation rate of monomers, *k_dim_* is the rate of dimerisation in the cytosol, and *α* is the fraction of monomers that become dimers during co-translation dimerisation (which will dissolve as dimers in the cytosol).

**Figure 3:**
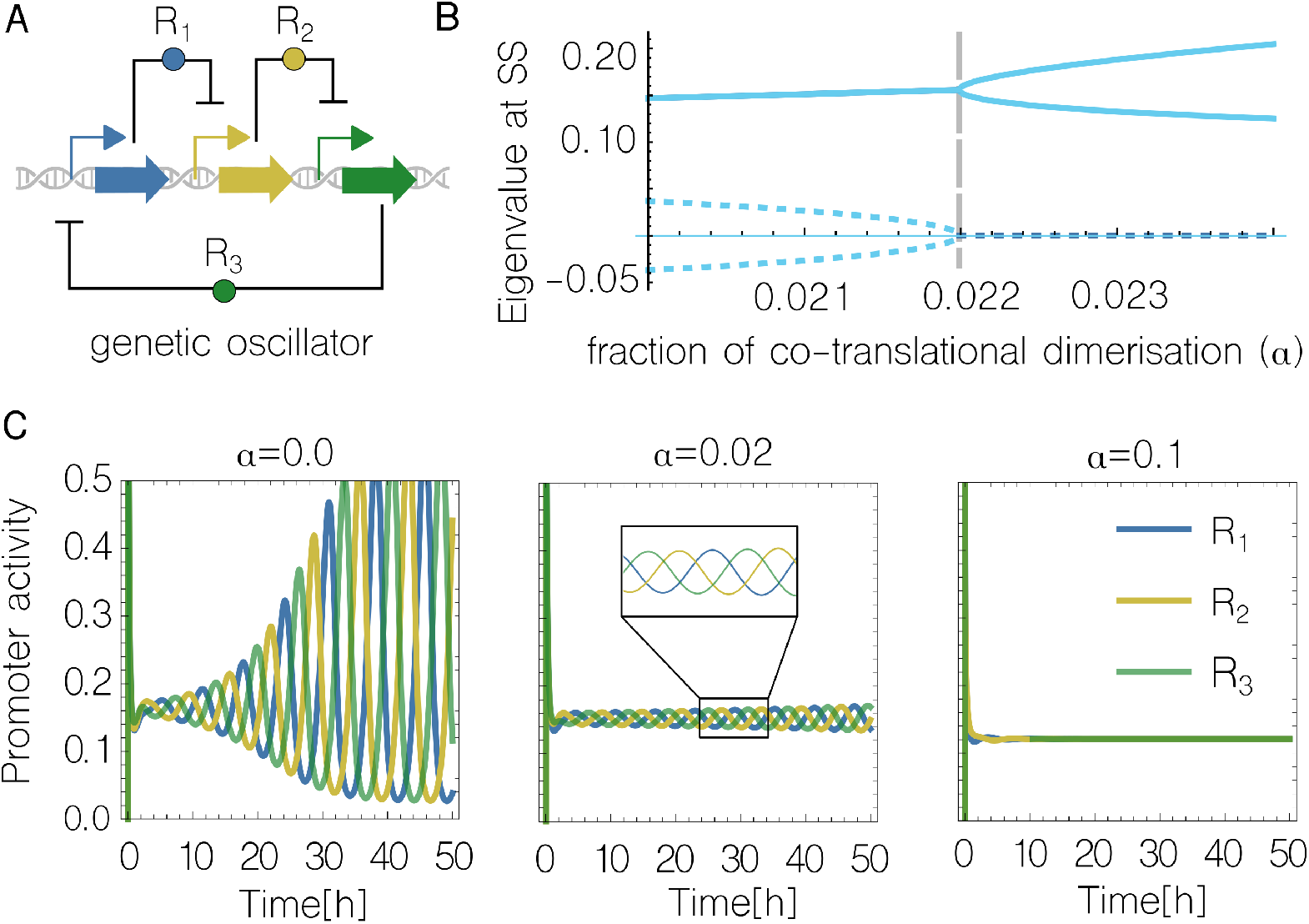
Impact of co-translational dimerisation (*α*) on a genetic oscillator. **A** Diagram of a three-component genetic oscillator (as in [45])—each repressor protein (R) inhibits the expression of its successor. As a result, the concentration of each repressor oscillates in time. **B** Eigenvalue analysis, which shows bifurcation point around *α* = 0.022. This suggests that the emergence of oscillations is highly dependent on the balance between co- and post-translational dimerisation. Solid and dotted lines are the real and imaginary values of the eigenvalues in the jacobian at equilibrium, respectively. **C** Time-course simulations of the genetic oscillator at different values of *α*. As shown, reliable oscillations are lost for values indicated in **B**.

Secondly, the ODE that calculate repressor dimers (i.e., the fully active protein) is given by:

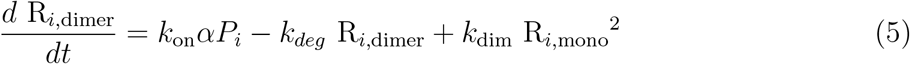

where *R_dimer_* is the dimerised repressor (note that in this model all three repressors of the system are assumed to be dimers in its final form). Lastly, the next ODE calculates the fraction of promoter *P* that is active (i.e., it is not being repressed by any R):

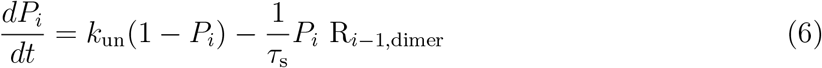

where *k_un_* is the unbinding rate of any dimer from its cognate promoter *P*, and 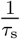 is the binding rate of the same two components. Note that promoter number *i* is repressed by a repressor number *i* – 1, where the previous element to *i* = 1 is *i* = 3, since the three genetic elements (1,2,3) are arranged in a ring.

Figure 3 shows how the parameter *α* alters the nonlinear patterns of the system, to the point that the circuit stops oscillating if the balance between co-vs. post-translational dimerisation is beyond a bifurcation point (Figure 3B). This is, higher values of *α* lead to a non-oscillating steady state (Figure 3C). This bifurcation, beyond which the oscillatory behaviour vanishes is around *α* = 0.022, which indicates that, even if the majority of dimers are formed in the cystosol, the ones that form during translation can drastically affect circuit behaviour.

The new parameter *α* introduces complexity to the mathematical model in that it limits the range of parameter values that generate oscillations—it could be argued that it is more difficult to get reliable oscillations than without it. However, it offers several pathways to modify nonlinearity in-vivo, unlike traditional methods for nonlinear dynamics (e.g., Hill coefficients). Therefore, its predictive scope narrows the gap that goes from modelling results to in-vivo experimentation.

### 2.3 Case study: programmabnle nonlinearity in a genetic toggle switch

A genetic toggle switch [36] is a device build from two mutually inhibitory repressors that is able to flip between stable states (Figure 4), in which only one of the two proteins is at high expression while the other one is inhibited. The stability of the system, and also the number of stable states, depends intricately on nonlinear dynamics. Although there are examples of genetic switches where nonlinearity emerges from protein dilution [46, 47], the most common way of achieving nonlinear dynamics (and bistability) comes from transcription cooperativity— in which protein dimerisation plays a fundamental role.

**Figure 4:**
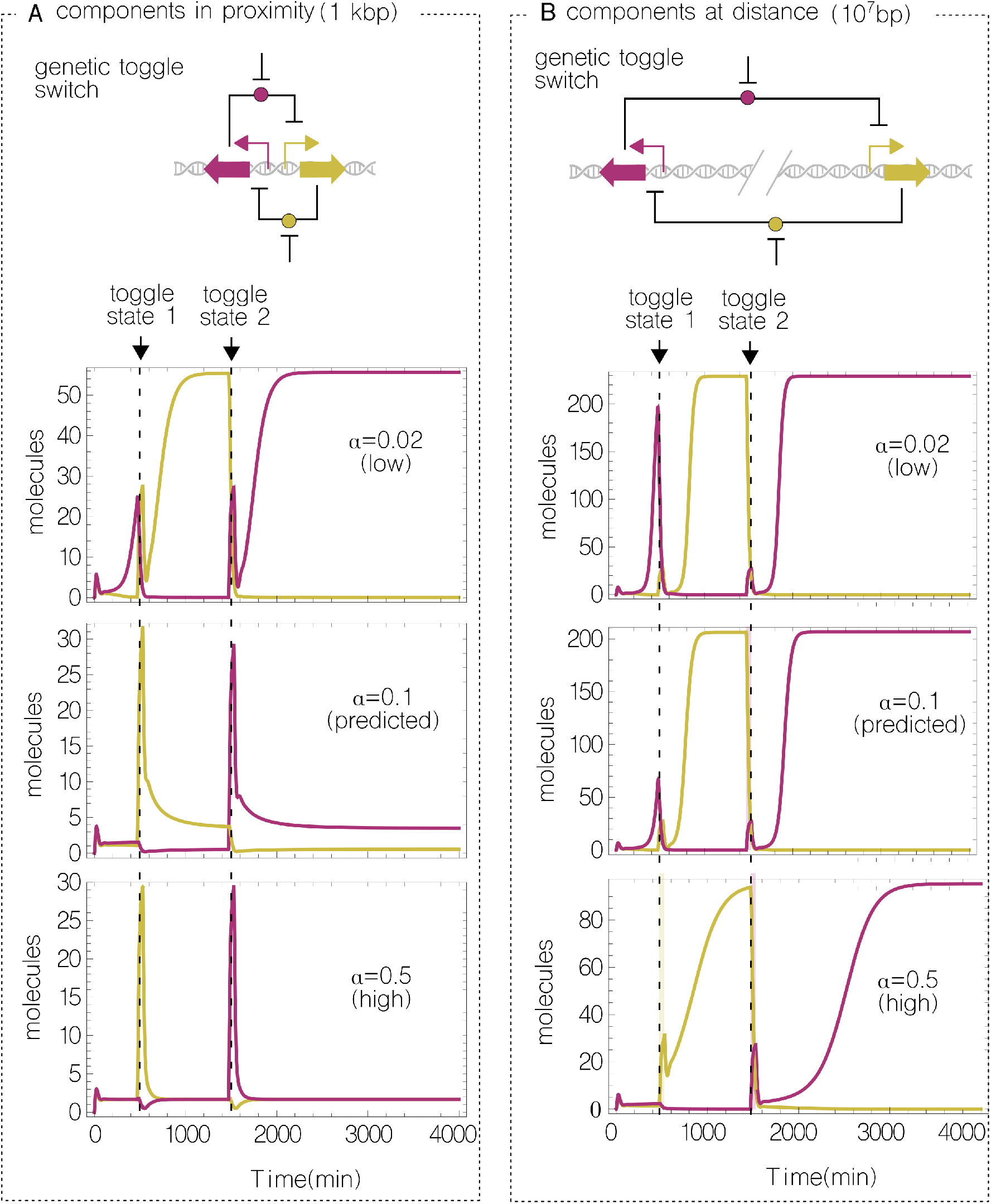
Impact of co-translational dimerisation and intergenic distance on a genetic toggle switch. **A** The two circuit modules are located in proximity (i.e., sharing the same chromosomal position). In this scenario, only in the case of *α* being low, there is enough nonlinearity to achieve bistability. **B** Genetic components are placed at distance (i.e., different chromosomal location). The physical separation forces proteins to *travel* from its source gene to its target promoter—a nonlinear process which counteracts the effect high *α* values. **A & B** The value of *α*=0.1 (tagged as *predicted*) is our theoretical approximation; up to now, and to the best of our knowledge, this value has not been experimentally obtained.

**Figure 5:**
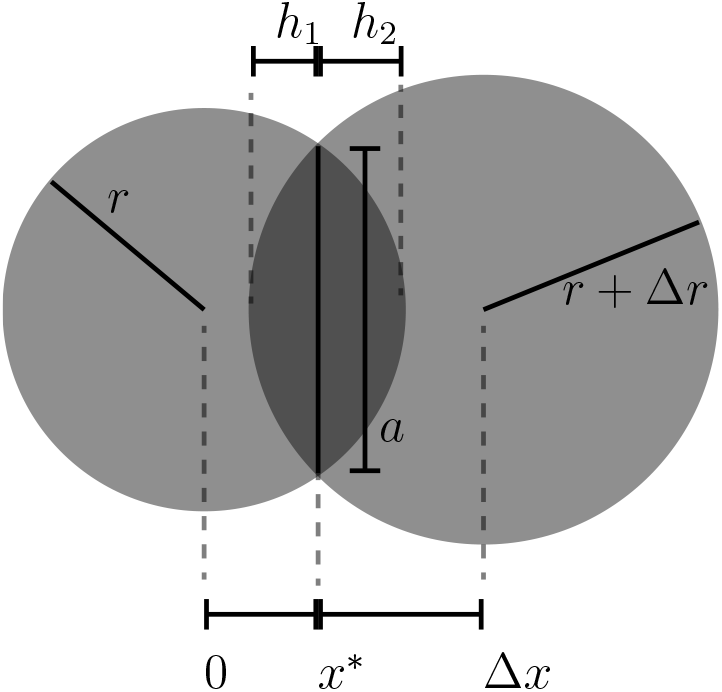
Calculation of the spherical overlap between partially formed monomers. At the centre of each sphere there is a ribosome bound to the RNA and actively translating. The radius (r and r+Δ*x*) define the leght of a monomer. Note that the dark grey overlapping section is composed of two spherical domes; for visualisation purposes, here we show the 2D analogous circular segments. The value of *h* is the height of the dome, and *a* the radius of its disk.

Our model for the toggle switch is specified by equations 6, 7 and 8, but with two (instead of three) repressor proteins (which are also assumed to be dimers). Furthermore, we added complexity to the model by considering the intracellular spatial distribution of genes, based on our recent work on spatiotemporal design [48, 35]. This feature helps differentiating between *local* and *global* repressors; this is, repressors that are in the proximity of, or far from, their encoding gene. The core message was that the distance a protein must “travel” from its source gene to its target promoter, modifies regulation in a predictable fashion. Since the differentiation between local and global proteins intersects with the model we introduce here, where proteins dimerise during translation (i.e., local) or in the cytosol during free diffusion (i.e., global), we analysed the genetic switch considering both dynamics.

Figure 4 shows results of simulating the switch with two different spatial setups, proximity (Figure 4A) and distance (Figure 4B), and three values for the fraction of co-translational dimerisation (*α*) in each case. Interestingly, the two spatial configurations show different performance for the same level of *α*, which implies that both phenomena (intergenic separation and protein dimerisation) are closely connected in shaping nonlinear dynamics.

When the genes of the switch are placed in proximity (Figure 4A), only a low value of α generates the sufficient nonlinearity for the system to show bistability. This is because the generation of dimers imposed by co-translational dimerisation is fully linear—so the fewer the better. As soon as this value increases, more dimerisation takes during translation, leading to system malfunction; although the switching still occurs to some extent (e.g., *α* = 0.5), output values are low, and the intrinsic stochasticity of biological systems would result in unstable states. However, moving genes far apart (Figure 4B) restored the function of the system, even in the case of *α* = 0.5 (a value we predict to be higher than physically plausible). By increasing the separation between the genes, the dimers formed near the source are not immediately available to bind to the target promoter, since this is now at a long distance. As a result, such linear process is now less relevant: co-translated dimers must “travel” from source to target resulting in a decreased promoter-binding rate, which, in turn, removes its involvement in total promoter binding events. Unlike the *proximity* scenario, where performance was only achieved at low values for α, the *distance* set up showed bistability at any value. This highlights the role played by intra-cellular distance for circuit design.

Altogether, simulations suggest that the necessary conditions for the emergence of bistability could be rationally designed or fine-tuned, and mapped into biological specifications such as ribosome binding sites (which impacts on α) or inter-genic separation.

## 3 Conclusion

The design of increasingly complex biological circuits in living cells is a major challenge. The lack of robust predictive modelling—which accurately foresee the performance of a design before implementation—threatens to undermine the success of the field. Although model-based design [49] is a common practice, accurate predictions are difficult to obtain since the internal workings of the cell are still based on unclear dynamics. Here, we focus on modelling protein dimerisation [33] within regulatory interactions, as a way predict and rationally design nonlinearity.

For protein dimerisation to take place, monomers must meet within the internal milieu of the cell. Although there are previous efforts that highlight the relevance of this issue [38, 39], the mathematical formalisation of the mechanistic details responsible for dimerisation has received little attention. Our model explicitly simulates co- and post-translational dimerisation, which occur during and after translation, respectively. While the former imposes a linear regime on the reaction, the latter boosts the emergence of nonlinear dynamics. By controlling the fraction of each dimerisation type, the model suggests routes for fine-tuning nonlinearity in vivo to fit pre-defined functional requirements. Moreover, the model was coupled to a previous model on spatiotemporal design that accounts for the diffusion of molecules within the volume of the cell [48]. Simulations suggested that intergenic distance (i.e., physical distance between two inter-regulated genes) plays a fundamental role in shaping dimerisation dynamics. Specifically, the linear dynamics imposed by co-translational dimerisation are less important if the two interacting genes are far apart; and that distance can be adjusted in-vivo by assigning different chromosomal locations to circuit components. Since nonlinear dynamics are at the core of biological [30] and computing [32] systems, its rational design would enable the construction of biological circuits with enhanced information-processing capabilities. The model presented here paves the way towards that goal.

Simulation results suggest a number of strategies on how to implement control over nonlinearity in-vivo. For instance, [i] protein length [50] (adding tags or extra amino acids), [ii] choice of RBS [51] (stronger RBS will result in closer ribosomes on the mRNA), or [iii] ribosome availability [52] (influencing binding rates)—let alone the modification of intergenic distance [35]. Future work on this line will focus on obtaining experimental measurements of the impact of co-translation based on the aforementioned implementation strategies. These will results in identifying the extent to which nonlinear dynamics can be adjusted on-demand. Furthermore, future theoretical work will aim at studying whether co-translational oligo-merisation could be modelled with a similar approach. Our model accounts for first-order interactions (i.e., monomers with monomers) but many functional transcription factors are, in fact, dimers of dimers. While the model, as is, can simulate the post-translational dimerisation of cotranslated dimers (thus representing regulators such as LacI, which is a tetramer), it cannot describe the formation of oligomers *during* translation. Although it is unlikely to play a major role on nonlinearity, this possibility will be targeted in future research on this line.

As precision is increasingly required from synthetic constructs, we advocate for the rational design of nonlinearity—a fundamental feature of biological systems. The model introduced here demonstrated the in-principle feasibility of it, and provides baseline information for the future in-vivo implementation.

## 4 Methods

### 4.1 Modelling the distance between ribosomes

In the model presented here, the distance between two consecutive ribosomes (Δ*x*) is determined by [i] the rate of translation and [ii] the difference in arrival time. This value is calculated as follows:

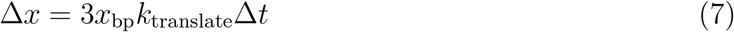

where *x*_bp_ is the width of a base pair, ca. 3*Å* [53], and the factor of 3 comes from one amino acid being encoded by 3 bp.

This difference in arival time (Δ*t*) also leads to a difference in length (Δ*r*) of partially translated proteins (or monomers). The calculation of the latter is captured by the next equation:

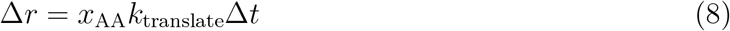

where *x_AA_* is the width of a contour length of an amino acid, also ca. 3 *Å* [54].

The model assumes this distance is constant until completing translation, which implies that the separation between ribosomes depends only on their binding times, not on sliding differences along the RNA.

### 4.2 Calculation of overlapping spheres

To calculate the overlap between the volumes of partially formed monomers we focus on a two consecutive ribosome scenario, which can be generalised to the ensemble of all ribosomes. In what follows, we show the derivation of equation 2 (i.e., overlaping fraction).

Here, we assume a system in which the two consecutive ribosomes have just initiated translation and are physically separated by a distance of Δ*x*. The radius described by the partially formed monomers is *r*_1_ and *r*_2_ respectively. In this scenario, the spheres are then described by:

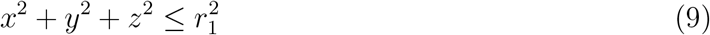

and

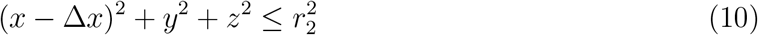

Using these equations, the edges of the spheres meet at a circle in the yz plane at location *x* determined by:

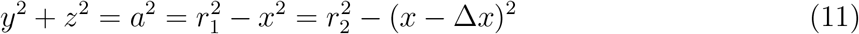

Which can be solved for *x*^*^:

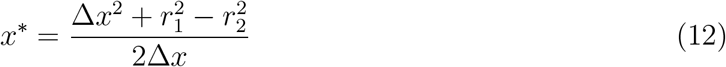

The overlapping volume can be described as a combination of two spherical domes (5) with the disk as described by equation 11 as the base. The volume of each dome is determined by:

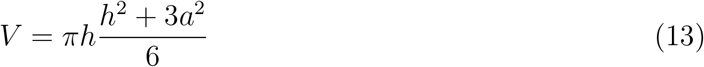

where *a* is the radius of the disk and *h* the height of the dome. The value of *h* differs between the two domes and is *r*_1_ – *x*^*^ and *r*_2_ – (Δ*x*) + *x*^*^ for the ribosme at the origin and the ribosome shifted by Δ*x* respectively. Upon assumption that the translation rate is constant the length of the transcripts of consecutive ribosomes is *r*_1_ = *r* and *r*_2_ = *r* + Δ*r*, with Δ*r* as determined by equation 8. The ribosome arrival time also determines the distance between the ribosomes (Δ*x*) via equation 7.

### 4.3 Code availability

All time-simulations, bifurcation analysis, derivations and data-visualisation were done in Wolfram Mathematica 12.0. Notebooks are availible at https://github.com/rstoof/TranslationMediatedDimerisation

## Acknowledgments

The authors acknowledge the SynBio3D project of the UK Engineering and Physical Sciences Research Council (EP/R019002/1) and the European CSA on biological standardization BIOROBOOST (EU grant number 820699). Lewis Grozinger is acknowledged for helpful discussions.

